# Genetic diversity and demographic history of the leopard seal: a Southern Ocean top predator

**DOI:** 10.1101/2023.04.05.535769

**Authors:** Arona N. Bender, Douglas J. Krause, Michael E. Goebel, Joseph I. Hoffman, Eric A. Lewallen, A. Bonin Carolina

## Abstract

Leopard seals (*Hydrurga leptonyx*) are top predators that can exert substantial top-down control of their Antarctic prey species. However, population trends and genetic diversity of leopard seals remain understudied, limiting our understanding of their ecological role. We investigated the genetic diversity, effective population size and demographic history of leopard seals to provide fundamental data that contextualizes their predatory influence on Antarctic ecosystems. Ninety leopard seals were sampled in the northern Antarctic Peninsula, during the austral summers of 2008–2019 and a 405bp region of the mitochondrial control region was sequenced for each individual. We uncovered moderate levels of nucleotide (π = 0.013) and haplotype (Hd = 0.96) diversity, and the effective population size was estimated at around 24,000 individuals (N_E_ = 24,376; range 16,876 – 33,126). Consistent with findings from other ice-breeding pinnipeds, Bayesian skyline analysis also revealed evidence for population expansion during the last glacial maximum, suggesting that historical population growth may have been boosted by an increase in the abundance of sea ice. Although leopard seals can be found in warmer, sub-Antarctic locations, the species’ core habitat is centered around the Antarctic, making it inherently vulnerable to the loss of sea ice habitat due to climate change. Therefore, detailed assessments of past and present leopard seal population trends are needed to inform policies for Antarctic ecosystems.

## Introduction

Marine mammals are increasingly being impacted by human-induced climate change and many polar species may not be able to respond at the rate required for their long-term survival [1, 2]. However, species-specific responses to changing environments are far from uniform, even within the same ecosystem (e.g., Arctic [3]). Surprisingly, population trends remain unknown for several conspicuous polar species, including top predators such as leopard seals, *Hydrurga leptonyx* [4]. Direct observations of this species in traditional surveys across the Antarctic are hindered by its broad and remote geographical distribution, low population density, and the adverse weather conditions that are commonly encountered in their main breeding habitats [5, 6].Therefore, indirect assessments, such as population genetic analyses, are invaluable for providing fundamental insights into the population dynamics and natural history of such species [7].

Contemporary neutral genetic variation retains valuable information that allows the inference of population sizes of past generations [8]. Specifically, the effective population size (N_E_) scales with the magnitude of genetic drift in an ideal population at equilibrium [9]. Most importantly, fluctuations in this parameter can be used to detect changes in species abundance over time, which can reveal past responses to environmentally and anthropogenically induced changes in habitat availability. For several species of pinnipeds, population genetic studies have uncovered past demographic impacts of large-scale climate change events driven by the El Niño Southern Oscillation [10] and the potential effects of post-glacial ice retreat on population expansion, recolonization and population structure [11-13]. These studies are fundamental for predicting how species will respond to future climate perturbations and for identifying taxonomic units that deserve prioritized conservation status.

Explorations of large genetic datasets across pinniped lineages also highlight biological traits that influence genetic variation over time. Breeding habitat preference (land vs. ice) is an important determinant of genetic variation in pinnipeds [14, 15], as substantial losses of genetic variation due to commercial sealing have been detected in gregarious pinniped species that breed on land [14]. Although this cumulative work represents a notable effort in terms of gathering samples and data for multiple pinniped species, some species have not yet been investigated. This reflects the difficulty of gathering samples from species breeding in remote, fluid and vast habitats such as the Antarctic pack ice.

The Lobodontini tribe (Antarctic ice seals) includes four species: the Ross seal (*Ommatophoca rossii*), the crabeater seal (*Lobodon carcinophagus*), the Weddell seal (*Leptonychotes weddelli*), and the leopard seal (*Hydrurga leptonyx*). The Weddell seal is one of the best-studied pinniped species [16] and is, therefore, typically included in foundational Antarctic habitat models (e.g., [17]). Conversely, Ross and leopard seals remain among the least studied pinnipeds [18]. Leopard seals have a broad circumpolar Antarctic and sub-Antarctic distribution [19], and are particularly challenging to study due to their solitary nature, pelagic lifestyle and scattered distribution [6]. Several visual surveys to estimate the abundance of leopard seals have been attempted [20-24], but most have yielded estimates with a high degree of uncertainty (Table 1). For example, the most recent global survey of leopard seals had large confidence intervals (N= 35,500, 95% CI: 10,900 – 102,600 individuals) because of the difficulty of counting this elusive predator, which spends a significant portion of time underwater and rarely aggregates on ice [24]. Therefore, genetic approaches have great potential to aid in our basic understanding of leopard seal population dynamics. To date, published genetic analyses of leopard seals are limited [25, 26] and descriptions of key demographic parameters such as N_E_ are lacking.

**Table 1.** Population sizes estimated for leopard seals, *Hydrurga leptonyx*, in Antarctica based on ship and aerial surveys. All estimates are rounded to the nearest 100.

| Source | Spatial coverage | Study year(s) | Abundance |
| --- | --- | --- | --- |
| **Eklund and Atwood (1962)** | Ross Sea<br>68°–70°S; 166°–177°E<br>Indian Ocean<br>64°–65°S; 105°–112°E | 1956 – 1957 | 152,500 |
| **Erickson and Hanson (1990)** | Most of the circumpolar pack-ice | 1968/69 – 1982/83 | 296,500 |
| **Ainley (1985)** | Ross Sea<br>62°–78°S; 106°–170°W | 1976/77 – 1979/80 | 8,000 |
| **Southwell et al. (2012)** | Most of the circumpolar pack-ice | 1998 – 2000 | 35,500<br>(10,900 – 102,600) |

Descriptions of leopard seal demography are essential to understanding the dynamics of the Antarctic ecosystem. As top predators, this species exerts top-down control on mesopredator populations [27, 28] and can disproportionally impact ecosystems [29]. For example, leopard seals have consumed an estimated 69.3% of all Antarctic fur seal pups (*Arctocephalus gazella*) born annually at Cape Shirreff in the north Antarctic peninsula since 2010, contributing to the rapid collapse of this regional population [28]. Leopard seals can also be considered vulnerable to the indirect effects of fisheries, which can reduce the availability of lower trophic level prey, such as krill, to their mesopredator prey [30]. Therefore, knowledge of historical and current global leopard seal population abundance trends are critical for robust ecosystem-based models (e.g., [31]) which are key to ongoing conservation efforts in the Antarctic [32, 33]. Here, we utilize mitochondrial DNA (mtDNA) sequence data from 90 leopard seals to characterize neutral genetic diversity, estimate the effective population size, and reconstruct the recent demographic history of this species. These data not only provide fundamental knowledge relevant to understanding leopard seal biology, but also contribute important data for inclusion in expanded Antarctic ecosystem models that consider the potential for leopard seals to alter ecological processes and/or be threatened by ecological destabilization.

## Materials and methods

### Sampling

Leopard seal sampling was conducted at Cape Shirreff (62°27’30’’S; 60°47’17’’W), on the north coast of Livingston Island, South Shetland Islands. The Cape encompasses several seasonally, ice-free gravel and pebble beaches bordered on its southern boundary by a permanent ice cap. We sampled a total of 90 leopard seals during identification-tagging efforts and captures during the austral summers (December to March) of 2008-09 through 2018-19. We obtained 2-4 mm^3^ of skin from the hind flippers utilizing manual hole-punch pliers, or sterile biopsy punches. All tissue samples were preserved in 95% ethanol and stored at −20°C. Sample collection efforts were undertaken by the U.S. Antarctic Marine Living Resources Program and information about sampled seals (e.g., Tag ID) is provided in Supplementary Table S1.

### DNA extraction and amplification

Total genomic DNA was extracted using a commercial kit (DNeasy Tissue Kit, QIAGEN) according to the manufacturer’s protocol. We amplified a 465-base pair (bp) fragment of the mtDNA sequence of the control region (D-loop) using pinniped-conserved primers TDKD for forward (5′-CCTGAAGTAGGAACCAGATG-3′) [34] and L15926 for reverse (5′-TCAAAGCTTACACCAGTCTTGTAAACC-3′) [35] annealing. PCR amplifications were performed using 25μl reactions containing 1μl of each primer (10μM), 2μl of Bovine Serum Albumin (BSA; 20 mg/μl), 12.5μl of GO Taq green master mix (Promega) and 2μl of template DNA (average concentration = 30ng/μl). Reactions were cycled in a T-100 thermocycler (Bio-Rad) according to the following protocol: 1 min at 94°, then 35 cycles of 94° for 1 min, 50° for 1 min and 72° for 1 min with a final extension at 72° for 7 min. PCR products were visualized on a 2% agarose gel with SYBR safe DNA stain (Invitrogen) to assess amplicon quality. Successful PCR products were enzymatically purified, and most individuals were sequenced in both directions using an Applied Biosystems 3730xl. The resulting forward and reverse sequences were manually edited using Geneious Prime v. 2021.1.2.2 [36]. The sequence ends were trimmed to remove primer sequences and to assure that only high-quality calls (quality score > 85%) were included in downstream analyses. The post-trimmed sequence length was 405 bp and the resulting sequences were aligned using MUSCLE [37].

### Genetic diversity

The number of haplotypes (h), nucleotide diversity (π), and haplotype diversity (Hd) were assessed using DnaSP v.6.12.03 [38]. We also reconstructed a haplotype network using the Templeton, Crandall, and Sing method (TCS) [39, 40] within PopART (Population Analysis with Reticulate Trees) [41].

### Effective population size

The effective population size (N_E_) was calculated using a Bayesian Most Probable Estimate (MPE) of the theta (Θ) value generated by LAMARC v.2.1.9 [42]. This method uses a Markov chain Monte Carlo (MCMC) sampling technique to estimate population genetic parameters by sampling parameter values as well as genealogies. Female effective population size (N_EF_) was estimated using the adjusted equation *N*_*EF*_ = Θ/2*μ*, where *μ* represents the mutation rate per site per generation. Assuming the population has a 1:1 sex ratio, the maternally inherited mitochondrial estimates were doubled to calculate the total N_E_.

### Neutrality test

Deviations from neutrality (whether a population has had a significant history of expansion or contraction) were investigated via mismatch distribution analysis within Arlequin v. 3.5.2.2 [43]. The mismatch distribution depicts the observed number of pairwise nucleotide site differences between all of the sequences found in a group of samples [44, 45]. Mismatch distributions are typically unimodal and smooth for populations that have recently expanded, while multimodal and ragged distributions are typical of stable populations [45]. Arlequin tests for population expansion by fitting a model of sudden expansion and calculates Harpending’s raggedness index (r) [46], which provides a measure of the smoothness of the empirical mismatch distribution. Expanding populations are expected to have lower values of r, whereas higher values are indicative of a stable population [46]. Neutrality tests (Fu’s F*s* and Tajima’s D) were also performed in Arlequin to infer demographic histories by determining whether the leopard seal sequences deviated from neutrality. Significant negative values indicate recent population expansion, while significant positive values indicate stable or bottlenecked populations [47-49].

### Bayesian skyline plot analysis

To infer the timing and magnitude of past changes in population size, we implemented demographic reconstruction using a Bayesian skyline plot (BSP) analysis in BEAST v.1.10.4 [50]. BSP uses patterns of coalescence and assumes a single panmictic population to fit a demographic model to a sequence data set [51]. We analyzed the data set under the HKY substitution model [52] with a strict molecular clock, in line with previous studies of other pinniped species that utilized the same mtDNA region (e.g., [53, 54]). We then used a coalescent Bayesian Skyline tree prior with six groups under a piecewise-constant model. Other priors used in this analysis were kept as default values. For the mutation rate, we opted for a value derived for the most closely related species, the Weddell seal (for the same mtDNA region: 1.14 × 10-7 substitutions per site per year (s/s/y)) [54]. We first converted the mutation rate units to s/s/gen (substitutions per site per generation) by multiplying Younger’s et al.’s rate by the generation time (GT) of leopard seals, which is estimated at 14 years [55]. The resulting mutation rate of 1.60 × 10^−6^ (s/s/gen) was then used in our analyses. All BSP analyses were run for 75 million MCMC iterations with parameters logged every 10,000 steps, and the first 10% were discarded as burn-in. Tracer v. 1.7.2 [56] was used to visualize the skyline plot reconstruction over time.

## Results

### Genetic diversity

We identified 34 mtDNA haplotypes among 90 individuals (GenBank accession numbers OQ451774-OQ451802; Supplementary Table S1), nearly half of which (47.1%) were singletons. Among the haplotypes detected, five had been previously reported for the species (Supplementary Table S1). The most common haplotype was detected in 10% of the samples and the most divergent haplotypes were separated by as many as 23 mutational steps (Fig. 1). The sequences were characterized by moderate levels of both nucleotide and haplotype diversity (π = 0.013; Hd = 0.96; Table 2).

**Fig. 1.**
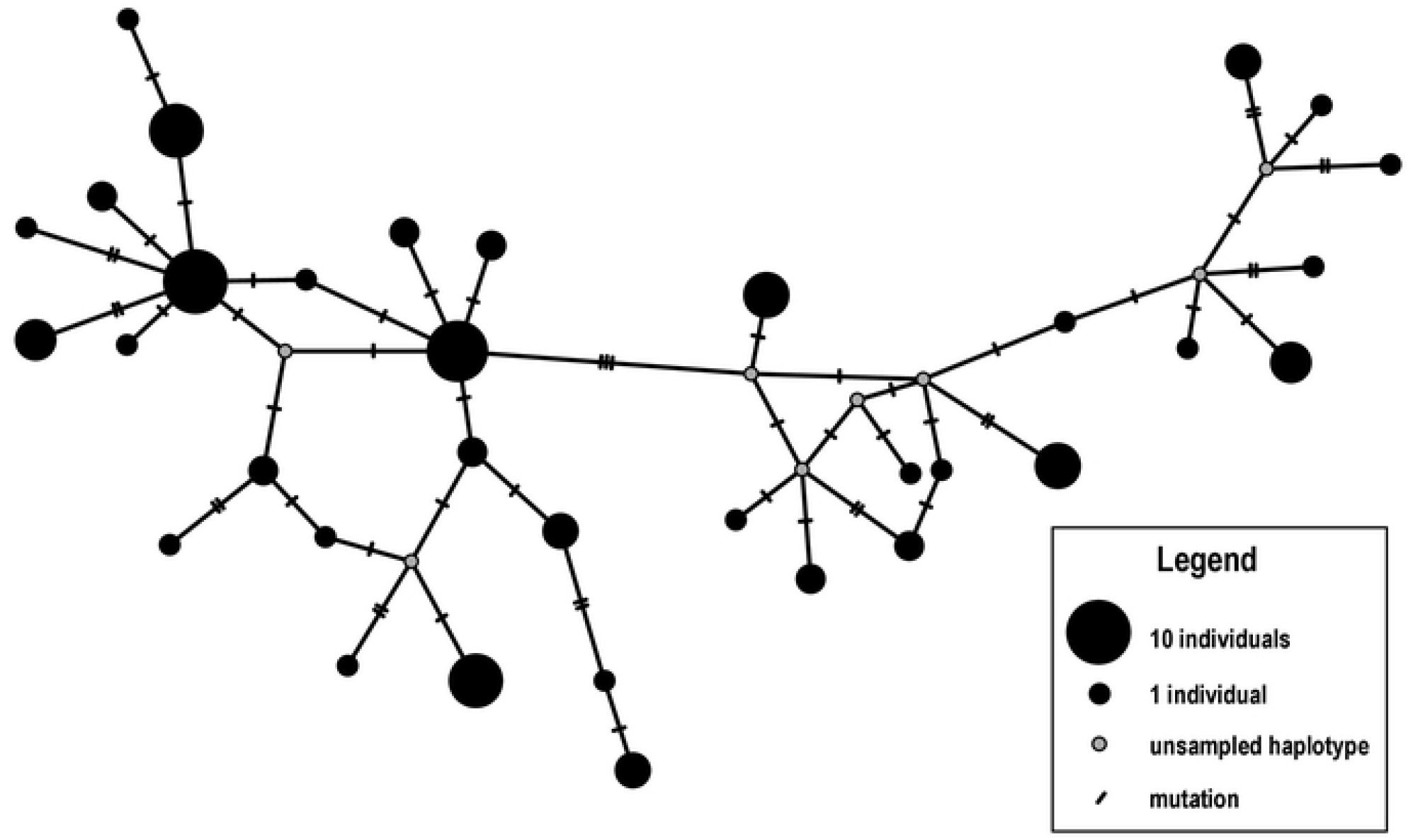
Haplotype network reconstruction of leopard seals, *Hydrurga leptonyx*, based on mtDNA sequencing data (n= 90; 405 bp mtDNA control region). This network was reconstructed via the TCS method [39, 40].

**Table 2.** Comparison of genetic diversity indices and the results of neutrality analyses among Lobodontini seals.

| Parameter | Leopard seal<br>(this study) | Weddell seal<br>(Curtis et al.<br>2009; 2011) | Crabeater seal<br>(Curtis et al.<br>2009; 2011) | Ross seal<br>(Curtis et al.<br>2009; 2011) |
| --- | --- | --- | --- | --- |
| **N** | 90 | 181 | 143 | 41 |
| **h** | 34 | 83 | 135 | 33 |
| **n singletons (%)** | 16 (47.1%) | 49 (59.0%) | 127 (94.1%) | 26 (78.8%) |
| **Hd** | 0.96 | 0.98 | 0.99 | 0.99 |
| **$\pi$** | 0.013 | 0.012 | 0.27 | 0.020 |
| **$\Theta$ (95 percentile)** | 0.039 (0.027–0.053) | 0.075 (0.055–0.085) | 0.576 (0.484–0.718) | 0.088 (0.060–0.132) |
| **Fu $F_s$ (P)** | -14.76 (0.00) | -24.99 (0.00) | -24.09 (0.00) | -20.13 (0.00) |
| **SSD (Mismatch P)** | 0.003 (0.58) | 0.002 (0.56) | 0.001 (0.82) | 0.06 (0.04) |
| **Raggedness (P)** | 0.008 (0.69) | 0.007 (0.84) | 0.002 (0.82) | 0.017 (0.05) |
| **$\tau$** | 3.77 | 5.74 | 12.59 | – |
| **D** | -0.297 (0.49) | – | – | – |

**Legend:** **N** = sample size; **h**= number of haplotypes; **n singletons**= number of singleton haplotypes; **Hd**= haplotype diversity; **$\pi$** = nucleotide diversity; **$\Theta$** = estimations of theta; **Fu $F_s$** = Fu's test of selective neutrality; **SSD**= sum of squared deviations, **Raggedness**= Harpending's raggedness index; **$\tau$** = peak of nucleotide pairwise distribution, **D**= Tajima's D test.

### Effective population size

We estimated Θ (theta) = 0.039 (0.027 – 0.053), which assuming a mutation rate (*μ*) for Weddell seals of 1.60×10^−6^ s/s/gen [54], yields an effective female population size estimate of 12,188 leopard seals (95% CI = 8,438 – 16,563). Assuming a 1:1 sex ratio, the total effective population size for the species was estimated as 24,376 individuals (95% CI = 16,876 – 33,126; Table 2).

### Demographic reconstruction

The observed mismatch distribution did not significantly depart from a unimodal shape (P > 0.05; Harpending’s raggedness index = 0.008, *P* = 0.69; peak of distribution (τ) = 3.77; Fig. 2a, Table 2). Additionally, neutrality tests yielded negative values (Fu’s F*s* = –14.76, P < 0.01; Tajima’s *D* = – 0.297, *P* = 0.49; Table 2), indicative of an excess of rare polymorphisms. Together, these results suggest a past leopard seal population expansion.

**Fig. 2.**
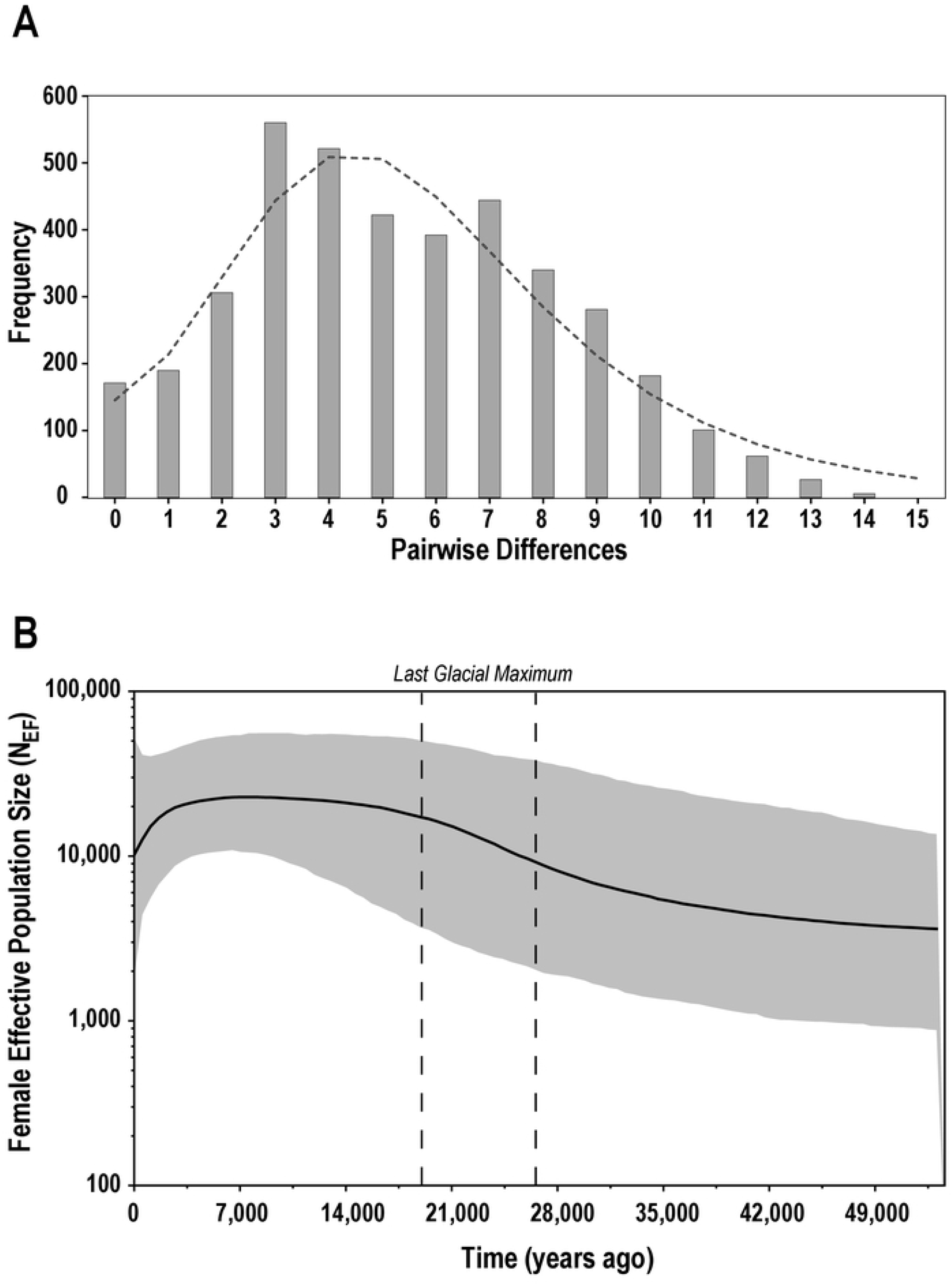
Demographic reconstruction of leopard seals, *Hydrurga leptonyx*, based on mtDNA sequencing data (n= 90; 405 bp). **A**. Mismatch distribution. Note that this distribution did not significantly depart from a unimodal shape (P> 0.05), rejecting constant population size; **B**. Bayesian Skyline Plot indicates timing of past expansion coincided with the Last Glacial Maximum (LGM; 26 – 19 KYA) [57].

To further investigate the timing of past population expansion, we reconstructed a historical timeline of female effective population size (N_EF_). Our analyses indicated a population that experienced an accelerated period of expansion during the Pleistocene Epoch, starting around 35,000 years before present (KYA; Fig 2b). This accelerated rate of expansion coincided with the last glacial maximum (LGM) around 26 – 19 KYA; a time of increased glaciation [57]. Subsequently, N_EF_ reached and remained at its highest levels from 13,000 – 6,000 KYA. This period was followed by a post-glacial population decline that continues until the present. The N_EF_ estimate from the BSP analysis was similar to our independent N_EF_ estimate based on Θ (BSP N_EF_ = 10,000 vs. LAMARC Θ N_EF_ = 12,000).

## Discussion

Due to the solitary nature of leopard seals and the vastness of their typical sea ice habitat, basic data on population trends are hampered by uncertainty. For this reason, leopard seals are not often incorporated into ecosystem models for the Southern Ocean, resulting in a knowledge gap regarding an important top predator. Here, we provide the first effective population size estimate for this species, based on a decadal sampling effort at Livingston Island, at the northern portion of the Antarctica peninsula. Our findings indicate that leopard seals have comparable levels of mitochondrial DNA diversity to their sister species, the Weddell seal. Furthermore, in line with other ice-breeding seal species [58, 59], historical population trends in this species appear to mirror the historical availability of sea ice, which was more extensive toward the end of the Pleistocene Epoch.

### Genetic diversity

Our estimates of nucleotide and haplotype diversity for leopard seals were comparable to findings in other phocid species. Haplotype diversity (0.96) aligned with reports for southern elephant seals (*Mirounga leonina*, Hd= 0.96, N= 203) [60] and Weddell seals (Hd = 0.98, N = 181) [58], but was lower than a previous assessment for leopard seals based on a much smaller number of samples (Hd = 0.99; N = 13) [26]. Haplotype diversity was moderate compared to hooded seals (*Cystophora cristata*; Hd ≈ 1.0, N = 123) [61] but very high compared to species that experienced strong bottlenecks such as northern elephant seals (*Mirounga angustirostris*; Hd = 0.41; N =185) [62]. Similarly, our estimate of nucleotide diversity for the leopard seal (π = 0.013) was within the expected range for phocids, although it was nearly half the value reported for the crabeater seal (π = 0.27), which has a population size estimated in the order of millions [21, 23]. Leopard seal nucleotide diversity was somewhat lower than the Ross seal (π = 0.02, N = 41), but remarkably close to the reported nucleotide diversity of Weddell seals (π = 0.012) [58]. Previous molecular data clearly support *Hydrurga* and *Leptonychotes* as sister taxa [63-66] and a proposed divergence circa 2.89 MYA [67] is consistent with the observation of similar genetic diversity parameters between these species.

### Effective population size vs. census size

The N_E_ estimate for leopard seals, 24,376 (95% CI: 16,876–33,126) is surprisingly high, considering the species’ trophic level and generation time. Our BSP analyses also indicates that N_E_ was historically large. We argue that this may be related to a generalist foraging strategy that allows leopard seals to exploit a variety of prey. Despite being a top predator, recent studies have shown broad intra-specific variability with regard to diet, which is quite diverse and varies seasonally [68]. Variability is also observed in diving behavior [69], hunting strategies [70], the use of ice floes [71] and movements [29]. In fact, the traditional notion that leopard seals are exclusively found in the Antarctic has also recently been rejected [19]. Individuals are recorded year-round in the sub-Antarctic and more northerly locations [19, 71], but the contribution of the northernmost populations to the overall species population size is unknown, and the species’ core breeding habitat is still presumably associated with circumpolar pack ice.

It would be helpful to extrapolate our N_E_ estimate for leopard seals to the census size (N_C_), but this presents additional challenges. Recently, a comprehensive multi-species study of N_E_ in pinnipeds revealed a mean ratio of N_E_/N_C_ of 31% [15], which is consistent with N_E_/N_C_ ratios of between 20% and 30% reported for the brown bear (*Ursus arctos*): a terrestrial species that is better characterized than most marine mammals [72, 73]. Utilizing the mean N_E_/N_C_ ratio of 31% [15], the contemporary leopard seal census population size (N_C_) can be inferred at 78,632 (95% CI: 54,438 – 106,868). In this context, the most recent circumpolar survey of leopard seals estimated N_C_ at 35,500 [95% CI = 10,900 – 102,600; 24] appears biased due to undercounting, but the 95% confidence interval nevertheless overlaps with our rough N_C_ estimate.

Our extrapolation based on the N_E_/N_C_ ratio has a couple of key caveats. Although N_E_ and N_C_ are undeniably correlated, their ratio is species specific and can be as low as 10% [74]. Additionally, N_E_/N_C_ is influenced by the choice of genetic marker, so its interpretation relies on precise knowledge regarding the distribution of genetic variation across the genome [75]; its impact on N_E_ is not fully understood. Therefore, an additional estimation of N_E_ for leopard seals based on genome-wide data would be beneficial as has been demonstrated with Antarctic fur seals [76]. Despite this, our findings suggest that leopard seal visual surveys may undercount animals. Indeed, underwater passive acoustic surveys revealed a much higher mean density of leopard seals detected by passive acoustics compared to visual surveys in the Davis Sea: visual density = 0.006 leopard seals/km^2^ vs. acoustics density= 0.31 seals/km^2^ [6].

### Demographic history

Demographic reconstruction revealed a population expansion during the late Pleistocene epoch. This expansion likely began approximately 35 KYA, which is somewhat more recent than proposed population expansions of Weddell seals in west Antarctica [58]; in fact, the peak of unimodal distribution (τ) for leopard seals is the smallest among Lobodontini seals. Environmental conditions during the late Pleistocene varied dramatically, but seasonal sea ice was perennial and extended to 45°S [77, 78]. Curiously, although the expansion was underway during the LGM (26 – 19 KYA), it reached its peak after this time (6 – 13 KYA). This suggests that environmental conditions around that time increased the amount of available sea ice habitat for the leopard seals but their population expansion was also potentially related to ice-associated prey availability (e.g., increased numerical abundance and/or species diversity). For example, crabeater seals, considered part of the diet of leopard seals in Eastern Antarctica [e.g., 79], had a population expansion earlier in the Pleistocene [58, 59] consisting an additional resource to leopard seals.

The post-glacial population decline of leopard seals detected in our analyses may be a consequence of more recent declines in the availability of breeding habitat, particularly in west Antarctica. This ice loss started during the Holocene (11.7 KYA) [80] and concurrently drove the population expansion of other pinniped species that benefit from ice-free conditions for breeding such as southern elephant seals [reviewed by 81] and Antarctic fur seals [13]. However, the recent decline that is evident in our skyline plot might alternatively be an artifact of the hidden effect of population structure on the BSP method [82]. For this reason, BSP analyses of significantly structured populations require a pooled sample approach (e.g., pooling samples from each sub-population) [54]. However, the only population structure study on leopard seals, based on the microsatellite genotypes of individuals sampled from six Antarctic and sub-Antarctic locations, revealed only very low levels of population differentiation (FST = 0.001 (– 0.002, 0.006)) [25], which appears negligible in this context. Additional historical demography analyses of leopard seals incorporating expanded geographic sampling should help clarify this finding.

Understanding the historical demography of a given species is indispensable to predicting the effects of global climate change, because ecological plasticity is inherently linked to intraspecific genetic variation [83], which in turn is tightly correlated with N_E_. This is particularly true for the leopard seal, which unlike other Antarctic pinniped species (notably Antarctic fur seals and southern elephant seals), did not experience confounding direct anthropogenic impacts such as sealing on a large scale [58].

In conclusion, our study shows that despite variability in leopard seal behavior and their occurrence year-round in warmer sub-Antarctic locations, the expansion of ice habitat during the Pleistocene played a key role in the species past abundance, and sea ice availability is likely to continue shaping this species’ demography into the future. In this context, the leopard seal emerges as a key indicator species of climate change in the Southern Ocean and should as such be regarded as an important component of future habitat modeling efforts in the region.

## Acknowledgments

This project was funded by the National Oceanic and Atmospheric Administration, Office of Educational Partnership Program with Minority Serving Institutions award number NA16SEC4810007 and NA11SEC4810002 (scholarship and research funds to ANB). CAB is supported by NSF HRD # 2000211 and NSF OPP # 2146068. We thank the former and current AMLR directors, Drs. R.S. Holt and G. Watters for their support. We are also grateful to the Cape Shirreff field personnel for assistance with sample collection over the years. Dr. D. Gibson (Marine and Environmental Sciences Department chair) provided key support for this project at Hampton University.

## Supporting information captions

**Table S1**. Leopard seals, *Hydrurga leptonyx*, sequenced for mtDNA control region (405bp). All individuals were tagged and sampled at Cape Shirreff, Livingston Island by the U.S. Antarctic Marine Living Resources (U.S. AMLR). GenBank accession numbers OQ451774-OQ451802 correspond to new haplotypes detected in this study.

